# Electrophysiological signatures of anxiety in Parkinson’s disease

**DOI:** 10.1101/2023.04.25.538260

**Authors:** Sahar Yassine, Sourour Almarouk, Ute Gschwandtner, Manon Auffret, Mahmoud Hassan, Marc Verin, Peter Fuhr

## Abstract

Anxiety is a common non-motor symptom in Parkinson’s disease (PD) occurring in up to 31% of the patients and affecting their quality of life. Despite the high prevalence, anxiety symptoms in PD are often underdiagnosed and, therefore, undertreated. To date, functional and structural neuroimaging studies have contributed to our understanding of the motor and cognitive symptomatology of PD. Yet, the underlying pathophysiology of anxiety symptoms in PD remains largely unknown and studies on their neural correlates are missing. Here, we used resting state electroencephalography (RS-EEG) of 68 non-demented PD patients with or without clinically-defined anxiety and 25 healthy controls (HC) to assess spectral and functional connectivity fingerprints characterizing the PD-related anxiety. When comparing the brain activity of the PD anxious group (PD-A, N=18) to both PD non-anxious (PD-NA, N=50) and HC groups (N=25) at baseline, our results showed increased fronto-parietal delta power and decreased frontal beta power depicting the PD-A group. Results also revealed hyper-connectivity networks predominating in delta, theta and gamma bands against prominent hypo-connectivity networks in alpha and beta bands as network signatures of anxiety in PD where the frontal, temporal, limbic and insular lobes exhibited the majority of significant connections. Moreover, the revealed EEG-based electrophysiological signatures were strongly associated with the clinical scores of anxiety over the course of the disease. We believe that the identification of the electrophysiological correlates of anxiety in PD using EEG is conducive toward more accurate prognosis and diagnosis and can ultimately support the development of new therapeutics strategies.

## Introduction

Anxiety is a highly prevalent psychiatric comorbidity in Parkinson’s Disease (PD), affecting up to 31% of the patients^1^, which is three times more prevalent than the general elderly population^2^. It can emerge at any stage of the disease, and be present even during the prodromal stage^3,4^. The clinical presentation of this disorder can include various subtypes ^1,5,6^ such as General Anxiety Disorder, non-episodic and episodic anxiety, panic attacks, social phobia, which can worsen motor symptoms ^7–9^ and cognitive functioning ^10–13^ and decrease the quality of life of patients^14,15^. Moreover, anxiety in PD comorbid often with other psychiatric symptoms such as depression and apathy ^16,17^, and the extensive overlap in their relevant features has hindered their clinical dissociation^18^. As a result, anxiety in PD is often underdiagnosed^1,19^ and undertreated^20^ yet limited scientific attention has been given to understand its underlying pathophysiology.

Non-invasive neuroimaging techniques are increasingly used to investigate the neural mechanisms of anxiety in PD^17,21^. Positron emission tomography (PET) and anatomical magnetic resonance imaging (MRI) studies have associated the anxiety in PD with reduced metabolism and cortical thickness in several subcortical regions including the amygdala, as well as in the bilateral anterior cingulate and prefrontal cortex^21–25^. Using fMRI resting state studies, functional disruptions in emotional-related cortical and subcortical regions were reported to correlate with anxiety symptoms^21,26–28^.

Electroencephalography (EEG) has been growingly employed to uncover the neural correlates of complex neuropathologies, such as neuropsychiatric disorders^29,30^. Providing direct measures of the neural activity, EEG has proven to be a valuable, non-invasive and cost-effective tool for biomarkers development. To date, only one study has compared anxious and non-anxious PD patients using EEG, revealing frequency-related spectral and functional disruptions, mainly in the frontal cortex, that characterize the anxiety in PD^31^. Yet, the use of EEG in case-control longitudinal studies to assess the neural correlates of anxiety in PD is still missing.

Here, we used High-Density (HD)-EEG recordings to excerpt the electrophysiological signature of anxiety in PD by comparing the spectral patterns and functional networks of anxious PD patients (PD-A) to non-anxious PD patients (PD-NA) and healthy controls (HC). We quantified the spectral signature in terms of network signature in terms of lobes-interactions and highest degree regions. We also explored the relationship between the EEG signatures at baseline and the clinical scores assessing anxiety at three and five years.

## Materials and Methods

### Participants

The study population, described in our previous studies^32,33^, was composed of PD patients and healthy controls (HC) enrolled from the Movement Disorders Clinic of University Hospital of Basel (city of Basel, Switzerland) as a part of a longitudinal study approved by the local ethics committees (Ethikkommission beider Basel, Basel; Switzerland; EK 74/09). The diagnosis of PD was based on the United Kingdom Brain Bank criteria for idiopathic Parkinson’s disease^34^. To be included in the study, patients had to meet specific criteria including a Mini-Mental State Examination (MMSE) score of 24 or above, no previous history of vascular or demyelinating brain disease, and sufficient proficiency in the German language. All participants provided written informed consent and were fully informed of the nature of the study. Included patients underwent neurological, neuropsychological, neuropsychiatric and EEG examinations at baseline (BL) and follow-up after a mean interval of 3 years (3Y) and 5 years (5Y).

As we focused on anxiety in PD, only participants that presented anxiety assessments were included in this study. Accordingly, 68 non-demented PD patients (22 females, age : 66.4 ±8.3) and 25 HC (10 females, age: 66.4 ±4) were selected at BL. As for the 3Y follow-up, the sample size was set to 42 PD patients (14 females, age: 70.5 ±7.9) and 17 HC (9 females, age: 68.9 ±6). Finally at 5Y, only 29 PD patients (12 females, age: 71 ±7) and one healthy control presented anxiety assessments and were included only for the correlation analysis. A flowchart of the study (Figure S1) as well as the main demographic, clinical and neuropsychological characteristics of the main cohort (Tables S1) and analysis cohort (Table S2) are presented in the supplementary materials.

### Neurological, neuropsychological and neuropsychiatric evaluations

Basic neurological and comprehensive neuropsychological examinations were carried out in all the participants. Patients were evaluated on their regular dopaminergic medication (‘‘ON’’ state) and the use of antidepressant and anxiolytics treatments was reported. The global cognitive score was assessed using the Montreal cognitive assessment score^35^ (MoCA), and patients were classified as with or without mild cognitive impairment (MCI) according to the Movement Society Task Force Level II criterias described in Litvan et al.^36^. Depression was measured using the Beck Depressive Inventory, second edition^37^ (BDI-II, German version) and apathy was assessed based on the Apathy Evaluation Scale^38^ (AES, German version).

Anxiety symptoms were evaluated using the German version of the Beck Anxiety Inventory^39^ (BAI), a 21 items self-rating scale. Each item is evaluated on a four-point Likert scale ranging from 0 to 3 (e.g., not at all; a little; moderate; or many). The total score ranges from 0 to 63 with higher scores representing increased symptoms severity. Leentjens et al.^40^ have validated the use of BAI in PD. As a score higher than 13 has been identified to show clinically significant anxiety, this cut-off was considered to divide the PD patients into two groups: PD patients with clinically relevant anxiety PD-A (N=18) and PD patients without anxiety PD-NA (N=50).

### EEG acquisition and preprocessing

Resting state EEG data were recorded for all participants using a HD-EEG system with 256 channels (Netstation 300, EGI, Inc., Eugene, OR). Participants were asked to relax, close their eyes and stay awake while seated in a comfortable chair for 12 minutes. The sampling rate was set to 1000 Hz. The raw EEG data were segmented into epochs of 40 seconds each and the first epoch of each recording was discarded from the analysis. As described in our previous study^32^, epochs were preprocessed automatically using the open-source toolbox Automagic^41^. Briefly, signals are subjected to band-pass filtering between 1 and 45 Hz, followed by the electrooculography (EOG) regression on 17 frontal electrodes to eliminate ocular artifacts. This step reduces the final number of channels to 239 which are mapped to four lobes of interest: frontal, parietal, temporal and occipital (see Table S3 and Figure S2 of the supplementary materials). Subsequently, bad channels exhibiting high variance (higher than 20 μV) or amplitude exceeding ± 80 μV are identified and interpolated. Finally, the artefact-free epochs were sorted according to their quality metrics^32^ and only the best six were retained for the rest of the analysis.

### Power spectral analysis

The Welch method ^42^ was used to estimate the power spectrum of signals at the scalp level. It consisted of computing a modified periodogram using the Hamming window with 20 seconds duration and 50% overlap to obtain the absolute power spectral density (PSD). The relative power spectrum was then computed by normalising each value of the absolute power spectrum by the total sum of the powers at each frequency of the EEG broadband (1-45 Hz). A [239 × 45] relative power features at the scalp level were thus obtained and used for further analysis.

### Functional connectivity analysis

The functional brain networks were estimated using the source-connectivity method^43^. First, the inverse problem was solved to reconstruct the dynamics of the cortical brain sources: the EEG channels and the MRI template (ICBM152) were co-registered, a realistic head-model was built using the OpenMEEG^44^ toolbox, and the weighted Minimum Norm Estimate (wMNE) method^45^ was applied on the cortical signals. The obtained source signals were then averaged into the 210 regions of interest (ROIs) of the brainnetome atlas^46^, which are mapped into seven cortical lobes of interest: Prefrontal (PFC), Motor (Mot), Parietal (Par), Temporal (Tmp), Occipital (Occ), Limbic (Lmb) and insular (Ins). Their affiliation is presented in Table S4 of the supplementary materials. Afterwards, the phase synchrony between different ROIs was computed using the Phase Locking Value (PLV) method^47^ and the dynamic functional connectivity matrices were estimated for six different EEG frequency bands: delta (1-4 Hz), theta (4-8 Hz), alpha1 (8-10 Hz), alpha2 (10-13 Hz), beta (13-30 Hz) and gamma (30-45 Hz). Those matrices were ultimately averaged across time and trials and their 21945 unique connections [=210 × 209/2] in each frequency band were used for further analysis.

### Statistical analysis

The statistical differences in demographic and clinical characteristics between the PD-A, PD-NA and HC groups were examined using the one-way analysis of variance (ANOVA). The chi-square test (for the categorical variables) and the independent samples t-test (for the continuous variables) were applied to examine the difference between the PD-A and PD-NA groups. Covariates such as age, sex, education levels and variables that showed significant differences between groups were included in the subsequent analysis.

Our main objective was to compare EEG-based features of the PD-A group to both PD-NA and HC groups. To accomplish this three-group comparison, we employed a two-step statistical process. First, we used a permutation-based non-parametric analysis of covariance (Perm-ANCOVA) to examine statistical differences in the relative power spectrum [239 channels x 45 frequencies] and functional connectivity networks [21945 connections x 6 bands] of the three groups at BL. We used 1000 permutations to identify the first set of significant power/connectivity features (*p<0*.*05*). As we were interested in identifying the features that predominantly represent the PD-A group, we defined two conditions: the PD-A_high_ condition, where the power/connectivity values of the PD-A group were significantly higher than both the PD-NA and HC groups (PD-A > PD-NA & PD-A > HC), and the PD-A_low_ condition, where the power/connectivity values of the PD-A groups were significantly lower than both other groups (PD-A < PD-NA & PD-NA < HC). Next, the second step of the process involved applying a two-tailed between-groups Wilcoxon test (corrected for multiple comparisons, *p<0*.*0167*) on the previous set of statistically significant features. Significant features that meet one of the above conditions were subsequently retained and considered as electrophysiological signatures of anxiety in PD.

### Anxiety signature scores and correlation analysis

In order to quantify the electrophysiological signature of anxiety in PD, two separate signature scores were defined: the spectral signature score (SSS) and the network signature score (NSS). The SSS is delineated as the ratio between the power indexes (PI) of the two previously defined conditions: PD-A _high_ *→ PI*_*high*_ and PD-A _low_ *→ PI*_*low*_:

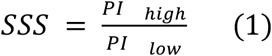

where *PI* is the mean relative power of the significant channels in the significant frequency slices and defined as:

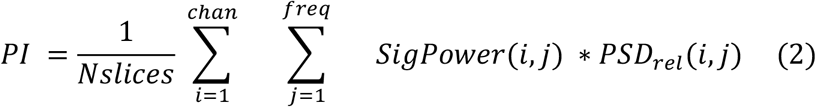

Where *SigPower* is a [239 × 45] binary matrix obtained from the statistical analysis representing the significant channels and their corresponding frequency slices, *PSD*_*rel*_ ± is the [239 × 45] matrix of the relative power features, *chan* is the total number of channels, *freq* is the total number of examined frequencies and *Nslices* is the total number of significant slices in *SigPower*.

Similarly, the NSS is defined as the ratio between the network indexes (NI) obtained from the significant edges of both conditions: PD-A _high_ *→ NI*_*high*_±and PD-A _low_ *→ NI*_*low*_:

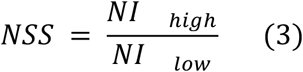

where *NI* is the mean connectivity of the significant edges (connections) in all frequency bands:

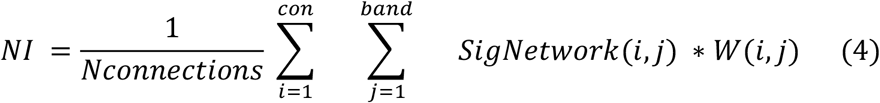

Where *SigNetwork* is a [21945 × 6] binary matrix obtained from the statistical analysis representing the significant connectivity features in each frequency band, *W* is the [21945 × 6] matrix containing the functional connectivity features, *con* is the total number of unique connections, *band* is the total number of EEG frequency bands and *Nconnections* is the total number of significant connections in *SigNetwork*.

A general mixed signature score (MSS) was also computed as the sum of the normalized SSS and NSS, representing thus both spectral and network signatures of anxiety in PD. Pearson’s correlation was used to examine the relationship between the electrophysiological signature scores (SSS/ NSS/ MSS) and the clinical anxiety score (BAI) not only at BL but also at 3Y and 5Y to assess their prediction capacity.

## Results

### Participant’s characteristics

Table 1 shows the demographic and clinical characteristics of the participants. No significant differences were found neither in the demographic features (age, sex and education) between all groups nor in the clinical assessments and the antiparkinsonian medication doses between the PD groups. Evidently, the BAI score was significantly discriminable between the three groups (p<0.0001). Also, both depression score (BDI-II) and apathy score (AES) presented a significant difference between groups (p<0.001) and significantly correlated with the BAI score. Therefore, they were both considered as covariates in the statistical analysis.

**Table 1.** Demographic and clinical characteristics of the three groups expressed as: mean (standard deviation). PD-A: PD patients with anxiety, PD-NA: PD patients without anxiety, HC: healthy controls, y: years, M/F: Male/Female, MoCA: Montreal Cognitive Assessment, MCI (Y/N): Mild Cognitive Impairment (yes/no), LEDD: Levodopa Equivalent Daily Dose, BAI: Beck Anxiety Inventory score, BDI-II: Beck Depression Inventory, second edition score, AES: Apathy Evaluation Scale. Significant p-values are marked in bold.

|  | PD-A<br>(N=18) | PD-NA<br>(N=50) | HC<br>(N=25) | P-value of<br>ANOVA | P-value of t-test<br>PD-A vs PD-NA |
| --- | --- | --- | --- | --- | --- |
| **Demographic** |  |  |  |  |  |
| Age (y) | 65.7 (8) | 66.6 (8.4) | 66.6 (4) | 0.89 | 0.68 |
| Sex (M/F) | 12/6 | 34/16 | 15/10 | 0.79 | 0.92 |
| Education (y) | 14.9 (3.7) | 14.7 (3.1) | 14.2 (2.9) | 0.74 | 0.78 |
| **Clinical** |  |  |  |  |  |
| Disease duration (y) | 4.9 (5.6) | 5.3 (5.1) | - | - | 0.79 |
| MoCA (/30) | 26 (2.8) | 26 (2.3) | 26.6 (2.7) | 0.63 | 0.99 |
| MCI (Y/N) | 5/13 | 17/33 | - | - | 0.73 |
| **Medication** |  |  |  |  |  |
| **LEDD<sup>48</sup> (mg/day)** | 616 (461) | 664 (470) | - | - | 0.71 |
| **Antidepressant (Y/N)** | 5/13 | 8/42 | - | - | 0.28 |
| **Anxiolytics (Y/N)** | 4/14 | 8/42 | - | - | 0.20 |
| **Neuropsychiatric tests** |  |  |  |  |  |
| **BAI (/63)** | 20.3 (7.2) | 6.2 (3.9) | 2.4 (3.2) | **<0.0001** | **<0.0001** |
| **BDI-II (/63)** | 11.2 (5.1) | 6.4 (4.1) | 2.6 (2.5) | **<0.0001** | **<0.001** |
| **AES (/63)** | 17.5 (10) | 6 (7) | 1(4) | **<0.001** | 0.29 |

### Spectral signature of anxiety in PD

The average relative spectral power over all EEG channels for the three groups is illustrated in Figure 1-A. Our statistical analysis on the overall [239 × 45] spectral features at BL allowed us to identify the spectral signature of anxiety in PD. This includes the EEG channels with their corresponding frequency slices where the PD-A group has either significantly higher or significantly lower spectral power than both the PD-NA and HC groups (PD-A_high_ and PD-A_low_ conditions). For the PD-A_high_ condition, results showed 20 significant channels with corresponding frequency slices mainly within the delta band (between 1 and 4 Hz). Those channels were presented notably in the parietal and frontal lobes. As for the PD-A_low_ condition, 11 channels mainly located within the frontal lobe and presenting significant frequency slices between 13 and 25 Hz (within the beta band) were revealed (Figure 1-B). The cortical topography of the relative spectral power observed in each group for the relevant frequency slices (delta and beta bands) of both conditions, along with the spatial distribution of the corresponding significant channels are illustrated in Figure 1-C.

**Figure 1.**
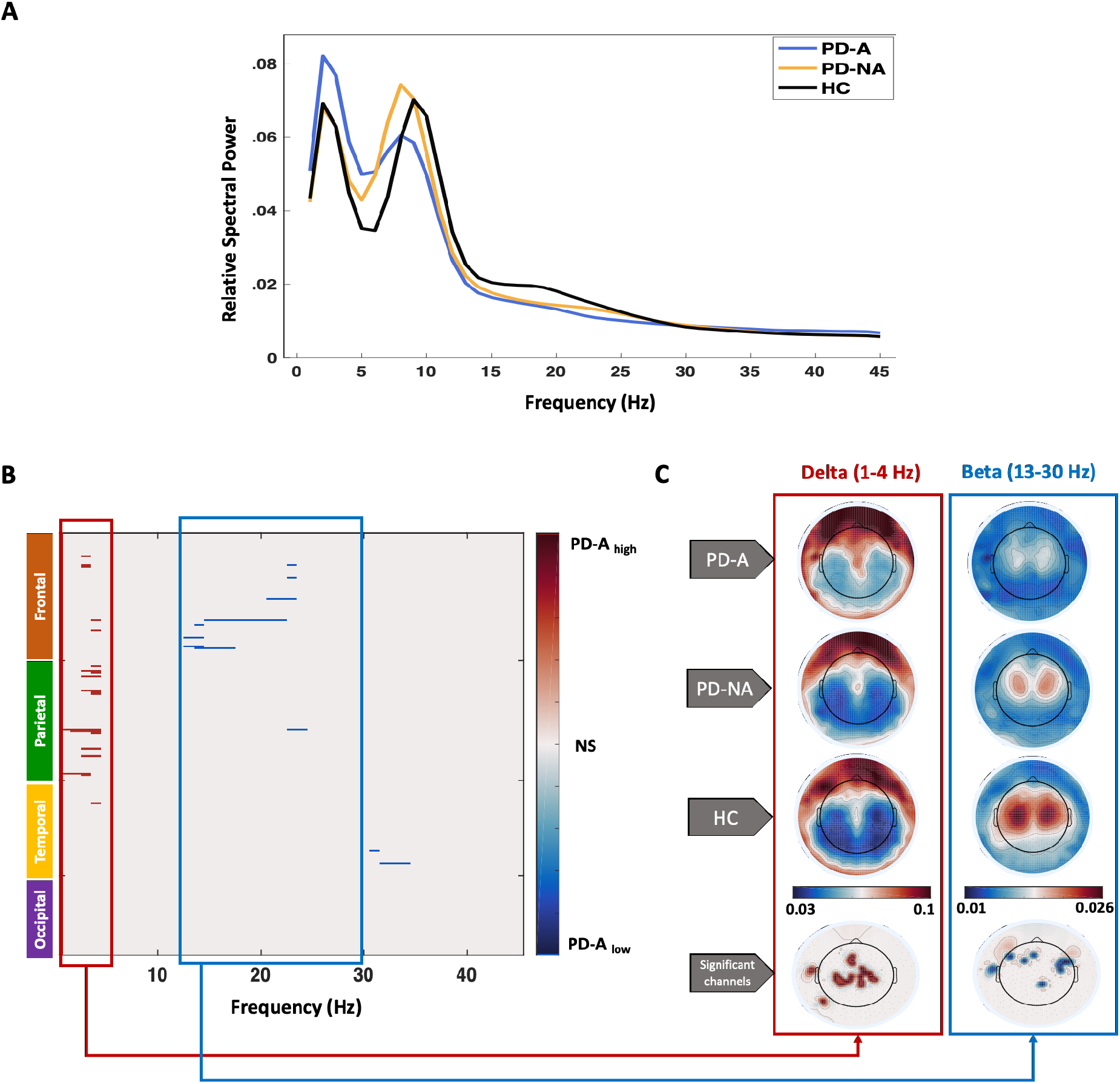
Spectral signature of anxiety in PD. **A)** the relative power spectra of the three groups: PD patients with anxiety (PD-A) and without anxiety (PD-NA) and healthy controls (HC). **B)** Significant channels and frequency slices of the PD-A_high_ (PD-A > PD-NA, HC) condition in red and PD-A_low_ (PD-A< PD-NA, HC) condition in blue. **C)** Cortical topography of the relative spectral power of the relevant frequency bands (delta in PD-A_high_ and beta in PD-A_low_) for the three groups and the corresponding spatial distribution of the significant channels (significant channels are marked in red for delta band and in blue for beta band). NS: no-significance.

### Spectral signature score

In order to appraise the spectral signature of anxiety in PD and associate it with clinical scores, we computed the SSS as the ratio between the average power of the significant channels/slices of the PD-A_high_ condition over the PD-A_low_ condition. Consequently, this resulted in investigating the spectral ratio between delta and beta bands. Results showed that the SSS of the PD-A group was significantly higher than both the PD-NA and HC groups (p<0.001, Bonferroni corrected, Figure 2-A). This SSS was significantly correlated with the BAI score (R=0.39, p<0.001) of the participants at BL (Figure 2-B). Additionally, when assessing the capacity of the SSS at BL in predicting the clinical scores of anxiety, results showed that the correlation remained significant with the BAI score at 3Y (R=0.41, p=0.011, Figure 2-C). A positive trend toward significance was also shown at 5Y (R=0.33, p=0.07, Figure 2-D).

**Figure 2.**
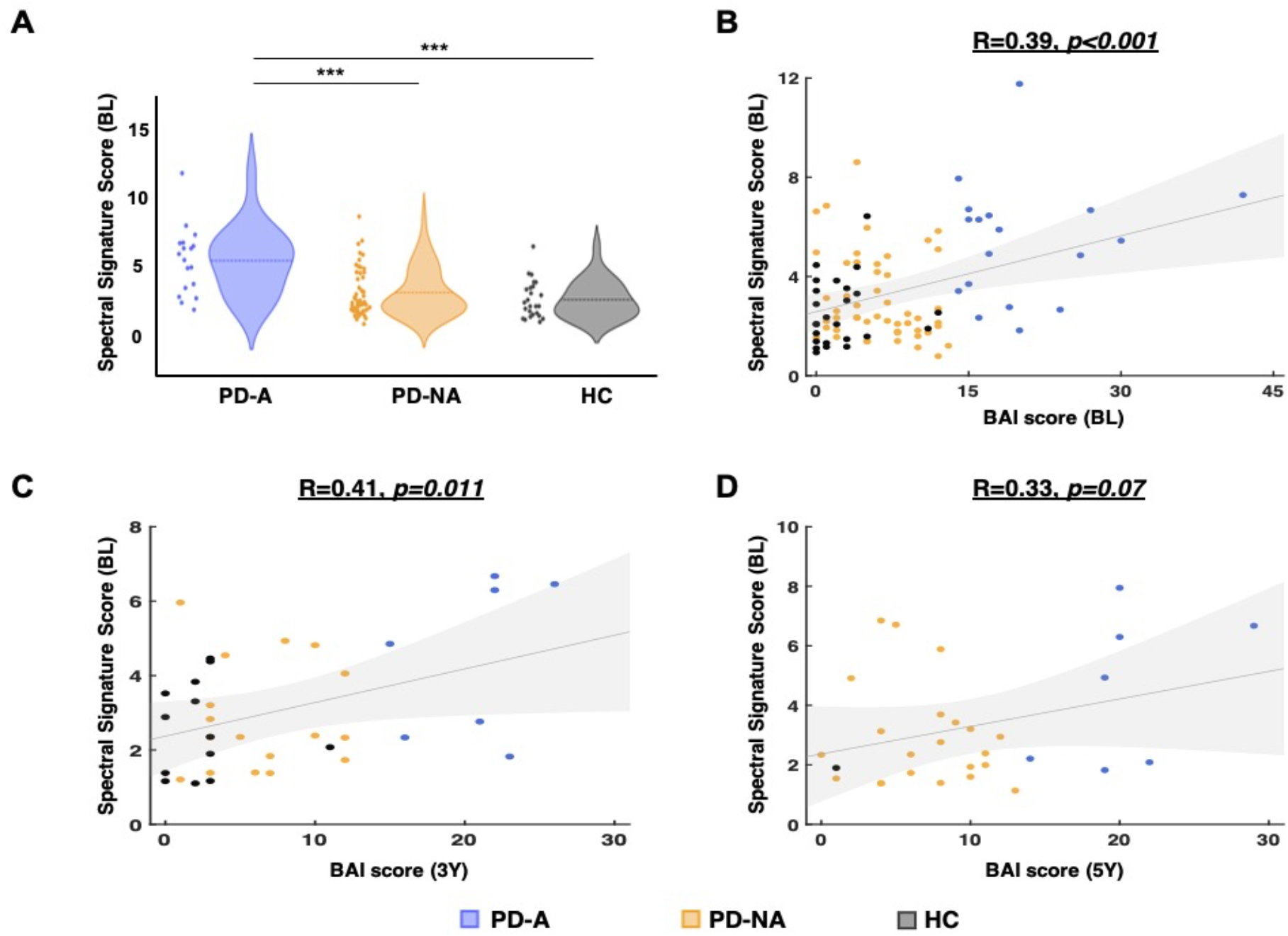
Spectral signature score (SSS) of anxiety in PD and its relationship with the BAI score. **A)** Distribution of the SSS between the three groups: PD patients with anxiety (PD-A), without anxiety (PD-NA) and healthy controls (HC). Relationship between the SSS at BL and the BAI score: **B)** at BL, **C)** at 3Y, **D)** at 5Y. *** p<0.001 (Bonferroni corrected for multiple comparisons).

### Network signature of anxiety in PD

Owing to uncovering the network signature of anxiety in PD, we repeated the same statistical analysis described above on the 21945 unique functional connectivity features of the six examined frequency bands. This resulted in identifying for each frequency band, a significant network of both hyper-connectivity edges (PD-A_high_ condition: where the connectivity in PD-A is significantly higher than in PD-NA and HC) and hypo-connectivity edges (PD-A_low_ condition: where the connectivity in PD-A is significantly lower than in PD-NA and HC).

Results showed that hyper-connectivity networks characterizing the PD-A group were dominant in delta, theta and gamma bands, while hypo-connectivity networks were more prevalent in alpha and beta bands (Figure 3-A). Further investigation of brian regions with the most number of connections (highest degree regions) in these significant networks revealed that regions within the temporal lobes were present in almost all bands. In particular the middle temporal gyrus (MTG) appeared in theta, alpha2 and beta bands. Additionally, the inferior frontal gyrus (IFG) was featured in networks of higher frequencies (alpha2, beta and gamma). Regions within the salience network (SAN) were among the most prevalent in theta (the caudodorsal region of the anterior cingulate gyrus (CG-cd)), alpha1 and gamma (the insula (INS)) (Figure 3-B).

**Figure 3.**
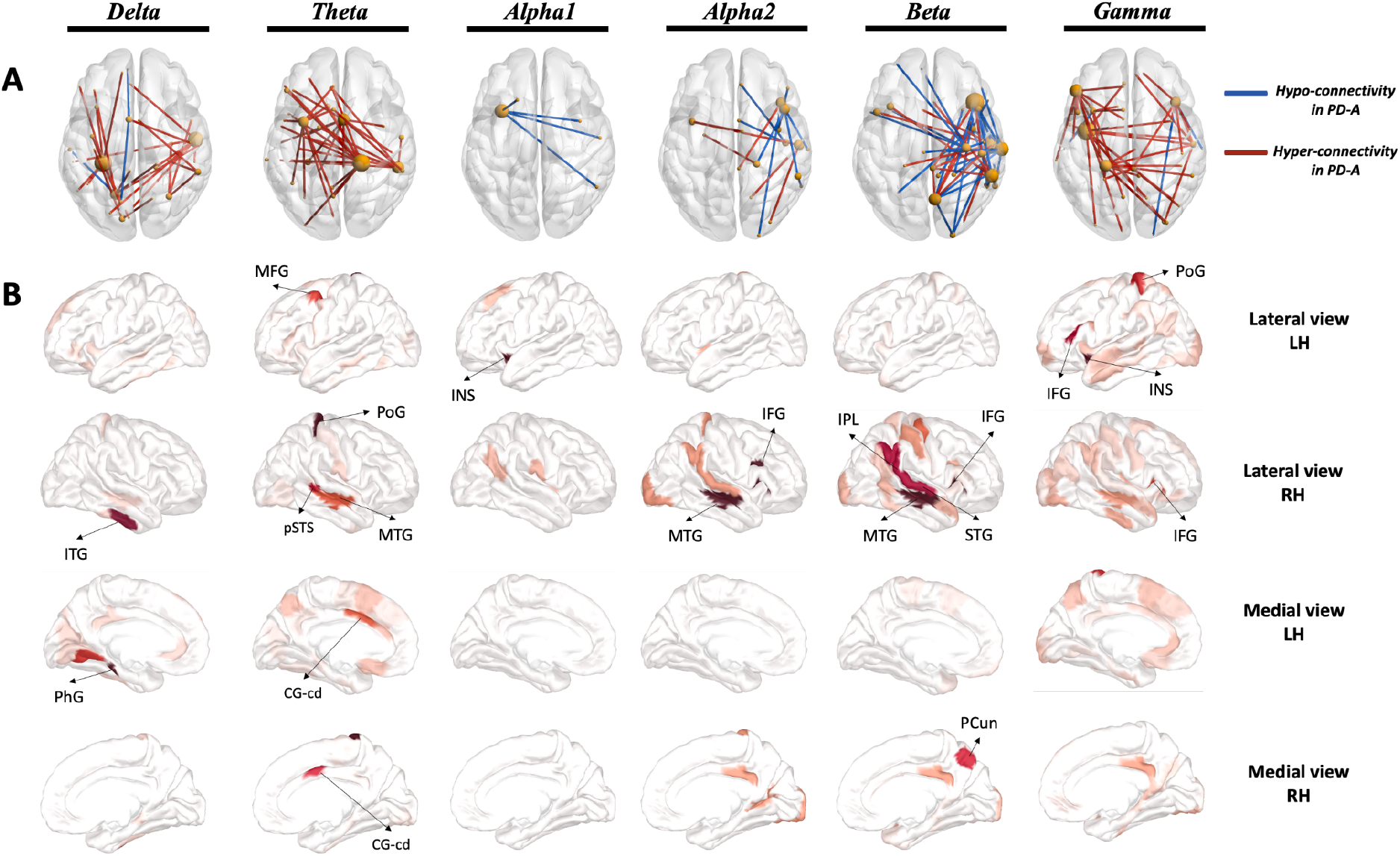
Network signature of anxiety in patients with PD. **A)** Significant networks of the different investigated frequency bands. The networks were thresholded for visualisation purposes. Edges presenting hyper-connectivity in PD-A are illustrated in red (PD-A_high_) and those presenting hypo-connectivity in PD-A are illustrated in blue (PD-A_low_). **B)** Highest degree regions (thresholded for visualisation purposes) represented with different views (lateral and medial) of the left hemisphere (LH) and right hemisphere (RH). ITG: Inferior Temporal Gyrus, PhG: Parahippocampal Gyrus, MFG: Middle Frontal Gyrus, PoG: Postcentral Gyrus, pSTS: posterior Superior Temporal Sulcus, MTG: Medial Temporal Gyrus, CG-cd: Cingulate Gyrus caudodorsal region, INS: Insula, IFG: Inferior Frontal Gyrus, IPL: Inferior Parietal Lobule, STG: Superior Temporal Gyrus, PCun: Precuneus.

Upon examining the interactions between the cortical lobes within these networks, we observed that the hyper-connectivity networks displayed dense functional connections primarily between the temporal, limbic and insular lobes. Specifically, the most prominent connections were temporo-temporal in delta, temporo-limbic in theta, motor-limbic in beta and insular-parietal in gamma bands. Regarding the hypo-connectivity networks, the insular lobe exhibited denser connections in the alpha band, with insular-parietal connections being the most dominant in alpha1 and insular-frontal connections prevailing in alpha2. Additionally, fronto-temporal hypo-connections were prevalent in beta bands. To illustrate these findings, circular and matrix plots displaying the interaction between the lobes of interest in the hypo/hyper connectivity networks across all bands are illustrated in Figure 4.

**Figure 4.**
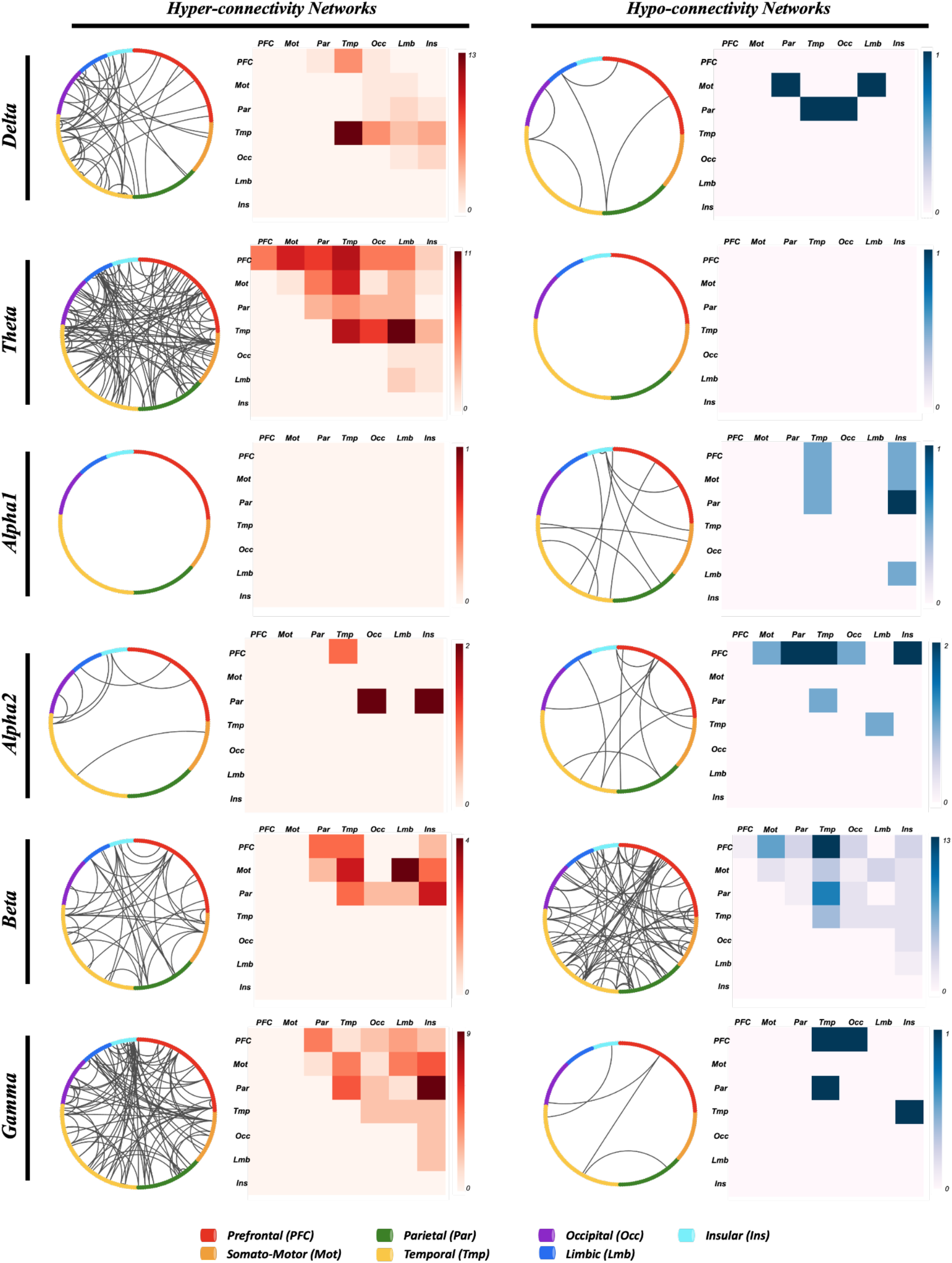
Representation of the network signature of anxiety in patients with PD. Circular plots (left) and matrix plot (right) of the significant networks Delta, Theta, Alpha1, Alpha2, Beta and Gamma frequency bands. Red and blue shades represent the number of connections in the hyper-connectivity networks and hypo-connectivity networks respectively.

### Network signature score

Following the analysis of the spectral signature, we also investigated the association between the network signature score (NSS) and the clinical evaluation of anxiety. This score represents the ratio between the average connectivity of the hyper-connectivity edges and that of the hypo-connectivity edges in all frequency bands. Results showed that this NSS was significantly higher in the PD-A group compared to both PD-NA and HC groups (p<0.001, Bonferroni corrected, Figure 5-A). The NSS at BL showed a strong correlation with the BAI score not only at BL (R=0.61, p<10^−10^, Figure 5-B) but also at 3Y (R=0.77, p<10^−7^, Figure 5-C). A positive trend toward significance was also shown after 5Y (R=0.33, p=0.07, Figure 5-D) demonstrating notable predictive ability.

**Figure 5.**
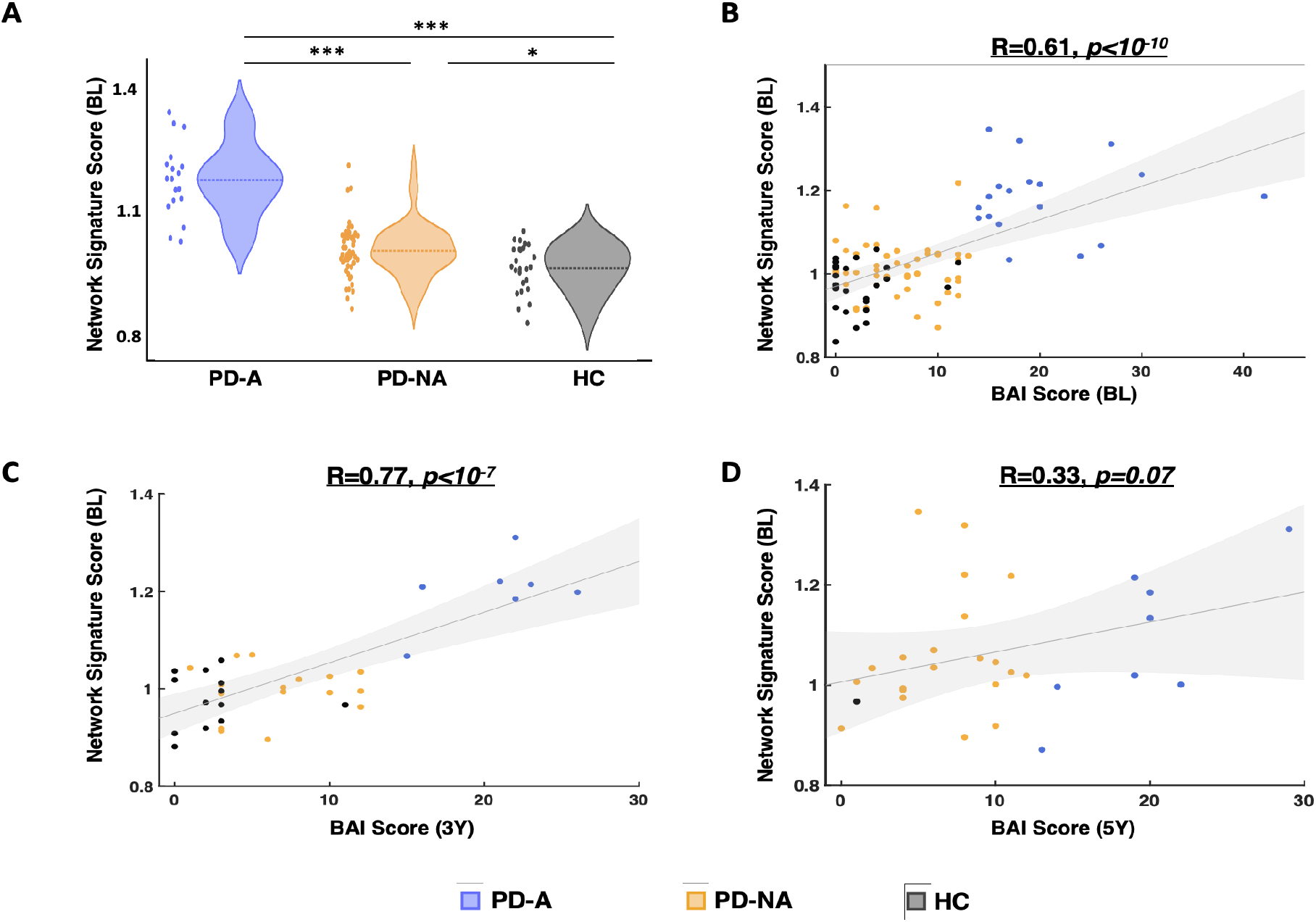
Network signature score (NSS) of anxiety in PD and its relationship with the BAI score. **A)** Distribution of the NSS between the three groups: PD patients with anxiety (PD-A), without anxiety (PD-NA) and healthy controls (HC). Relationship between the NSS at BL and BAI score: **B)** at BL, **C)** at 3Y, **D)** at 5Y. *** p<0.001, ** p<0.01, * p<0.05 (p-values are corrected using Bonferroni for multiple comparisons).

### General mixed signature score

The general mixed signature score, which combines both spectral and network signatures of anxiety in PD, demonstrated significant differences between the PD-A group and both PD-NA and HC groups (p<0.001, Bonferroni corrected, Figure 6-A). When examining its correlation with the clinical score of anxiety, the results showed that the MSS was strongly correlated with the BAI score not only at BL (R=0.52, p<10^−6^, Figure 6-B) but also at both follow-up examinations after 3Y (R=0.61, p<10^−4^, Figure 6-C) and 5Y (R=0.37, p<0.05, Figure 6-D).

**Figure 6.**
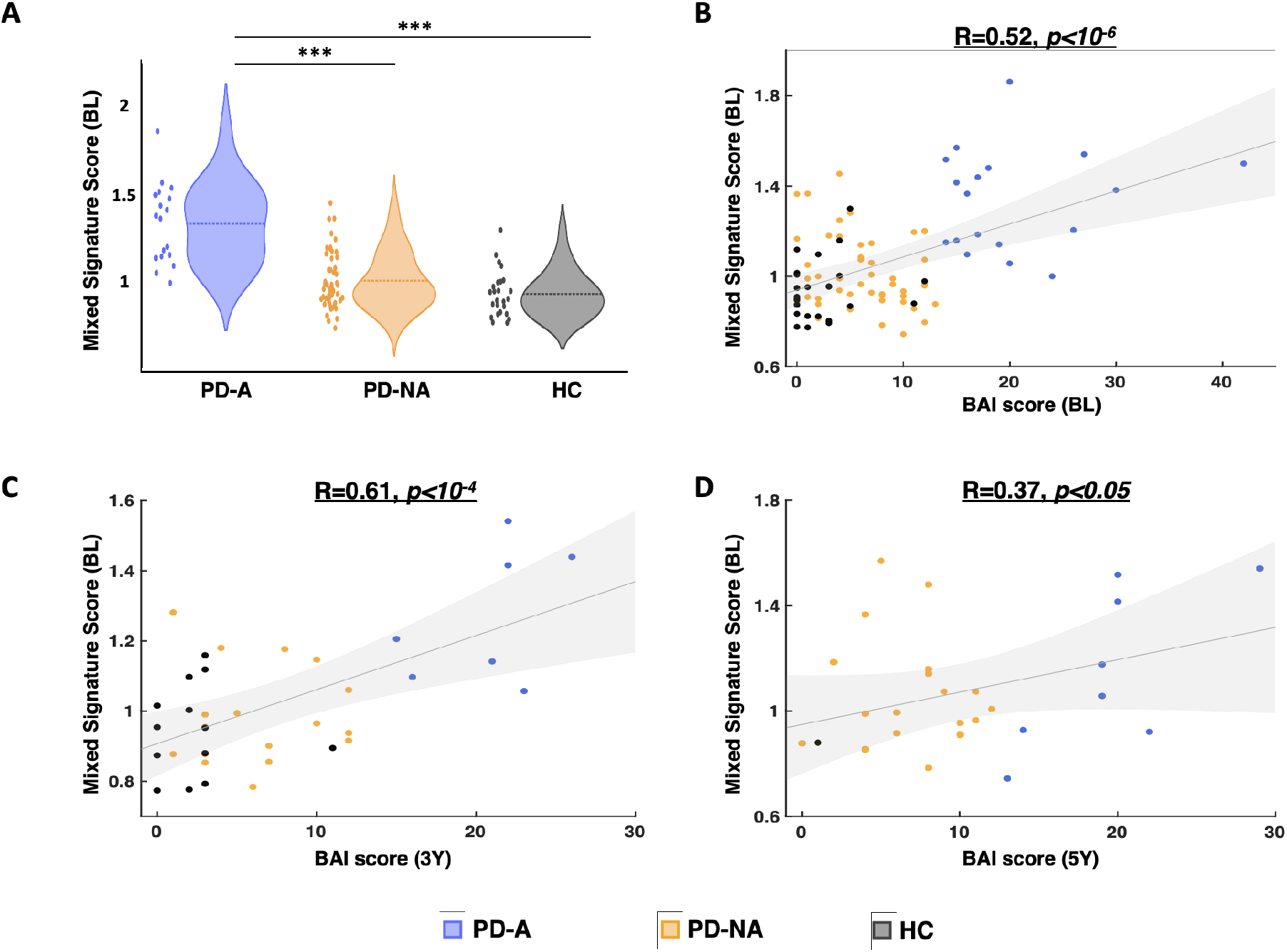
The general mixed signature score of anxiety in PD and its relationship with the BAI score. **A)** Distribution of the mixed signature score between the three groups: PD patients with anxiety (PD-A), without anxiety (PD-NA) and healthy controls (HC). Relationship between the mixed signature score at BL and BAI score: **B)** at BL, **C)** at 3Y, **D)** at 5Y. *** p<0.001 (p-values are corrected using Bonferroni for multiple comparisons).

## Discussion

In the present study, we aimed to identify the electrophysiological signatures of PD-related anxiety using resting state HD-EEG. While controlling the presence of other neuropsychiatric symptoms (depression and apathy), we showed that anxiety in PD is characterized by increased delta power -at the scalp level-in the frontal and parietal lobes as well as reduced beta power in the frontal lobe. Our functional connectivity analysis revealed that hyper-connectivity networks dominate in delta, theta and gamma bands while hypo-connectivity networks are more present in alpha and beta bands, with the frontal, temporal, limbic and insular lobes exhibiting the majority of significant connections. Electrophysiological scores (SSS/ NSS/ MSS) computed from the spectral/network signatures distinguish the PD-A group from both PD-NA and HC groups and are correlated with the clinical scores of anxiety at BL as well as at 3Y and 5Y, demonstrating predictive capacity.

Our spectral analysis at the channel-frequency level allowed for an accurate spatial-spectral mapping of the power features that characterize the PD-A group compared to both the PD-NA and HC groups. The increased power in delta and decreased power in low beta (13-20 Hz) are consistent with the global spectral patterns observed in the single previous EEG study that compared PD-A and PD-NA patients^31^. Our findings were also consistent with spectral patterns observed in anxious non-parkinsonian subjects. Increased delta power in frontal and parietal lobes was reported to characterize induced anxiety in obsessive compulsive-disorder patients^49^. Negative correlation between the powers of delta and beta bands in frontal regions was also shown in highly anxious healthy females performing a social task^50^. In addition, decreases in absolute and relative powers of slow and fast beta were observed in anxious adolescents^51^ and in patients with social phobia^52^. Nonetheless, positive delta-beta correlations and decreases in delta power have been also reported in social anxiety disorders but in studies with low-density EEG systems ^52,53^. Spatially, the frontal lobe was the most featured in our PD-anxiety spectral signature. Of interest, disruptions in the prefrontal cortex were consistently reported in neuroimaging studies, characterizing anxiety disorders not only in PD patients ^22,23,25,26,31^ but also in non-PD individuals ^50,52,54^.

Regarding the network signature of PD-related anxiety, we have demonstrated that hyper-connectivity networks were mostly dominant in delta, theta and gamma bands. Previous functional connectivity studies have associated increased severity of anxiety in PD patients with increased functional connectivity between cortical regions of the orbito-frontal cortex and both the inferior-middle temporal and parahippocampal gyri^28^ as well as between the insular lobe and both the prefrontal, and cingulate cortices^31^. These findings support the manifestation of the insula, the caudodorsal region of the cingulate gyrus, and the regions within the temporal and frontal lobes as well as their interactions as the most implicated in the hyper-connectivity networks of our results. Indeed, the insula along with the dorsal anterior cingulate (limbic) cortex and the medial prefrontal cortex are all parts of the fear/anxiety circuitry^55^ and activations and abnormalities in those regions have been consistently reported in different types of anxiety disorders in the general population ^56–59^ and in PD subjects^21,25,31^. This can be interpreted by the pivotal role of these core regions in processing fear, negative affect, worrisome thoughts and emotions ^60–62^. Additionally, hyperconnectivity between subcortical regions, mainly the amygdala and the putamen, and cortical regions of the fear/anxiety circuitry were also persistently associated with anxiety in PD^21,28,63^. While our analysis included only cortical regions, the dominance of the hyperconnectivity networks suggests a positive cortical-subcortical correlation between oscillations that stem from these regions.

Furthermore, we observed hypo-connectivity networks in alpha and beta bands, predominantly in the frontal and insular lobes. Consistent with our findings, previous research has shown that patterns of decreased connectivity within the frontal lobe are indicative of anxiety in PD patients ^25,28^. Moreover, functional dysconnectivity within and between the salience network, which involves mainly the insular lobe, has also been reported to reflect anxiety disorders in non-PD individuals^57,64–66^.

Importantly, our hypo/hyper-connectivity networks were also shown to be associated with the clinical traits of anxiety in all participants not only at baseline but also longitudinally after 3 years and 5 years. This association, validated also when combining both spectral and network signatures, can highlight the predictive capacity of our EEG-based markers of anxiety. However, despite this internal/longitudinal validation of our anxiety signature, external validation on an independent cohort is necessary for further endorsement.

Finally, some patients in both PD-A and PD-NA groups were under antidepressant and anxiolytic medications during EEG and neuropsychological assessments sessions. Here, we controlled for this issue by demonstrating that the anxiety and depression medication statuses did not differ significantly between PD groups. Besides, topographic EEG changes reported in generalized anxiety disorders during anxiety treatments^67,68^ suggested decreased spectral power of delta and alpha bands along with increased power of beta band^69–71^. Antidepressant medication^72^ has also been shown to reduce slow wave EEG activity and increase the power in alpha band^73,74^. Notably, these spectral patterns were not reported in our study to characterize the PD-A group, which included patients taking anxiolytics and antidepressants. Excluding these patients would have been an alternative solution in this study, however this would have reduced the sample size in the PD-A group by half and subsequently restricted our statistical analysis. Therefore, further research studying the neural correlates of anxiety in PD patients without anxiety/depression medications is still necessary for further validation.

## Conclusion

To summarize, this is the first case-control resting-state HD-EEG study to investigate the neural correlates of anxiety in PD. Our findings suggest that increased fronto-parietal delta power, decreased frontal beta power, and prevailed hyperconnectivity in several frequency bands are all EEG-based signatures of the PD-related anxiety. These signatures have the longitudinal predictive capacity for clinical outcomes. Identifiying such non-invasive markers may provide new perceptions into the development of advanced biomarkers. Further research could also establish resting-satte HD-EEG as a tool for more accurate prognosis and diagnosis of anxiety in PD and contribute in elevating the development of corresponding effective therapies.

