## Supplementary materials for "Electrophysiological signatures of anxiety in Parkinson’s disease"

**Table S1- Demographic, clinical and main neuropsychiatric characteristics of the main cohort longitudinally expressed as: mean (standard deviation).** y: years, M/F: Male/Female, MoCA: Montreal Cognitive Assessment, MCI (Y/N): Mild Cognitive Impairment (yes/no), UPDRS-III: Unified Parkinson's Disease Rating Scale-motor examination, LEDD: Levodopa Equivalent Daily Dose, BAI: Beck Anxiety Inventory score, BDI-II: Beck Depression Inventory, second edition score, AES: Apathy Evaluation Scale.

| | **Baseline** | | **3 years** | | **5 years** | |
|---|---|---|---|---|---|---|
| | **PD (N=77)** | **HC (N=32)** | **PD (N=45)** | **HC (N=21)** | **PD (N=44)** | **HC (N=3)** |
| ***Demographic*** | | | | | | |
| Age (y) | 66.2 (8.2) | 65.3 (5.6) | 70.9 (7.9) | 68.7 (4.9) | 71.7 (7.8) | 65.6 (4.1) |
| Sex (M/F) | 51/26 | 18/14 | 31/14 | 9/12 | 28/14 | 2/1 |
| Education (y) | 14.6 (3.2) | 13.8 (2.9) | 14.8 (3.1) | 13.6 (3.1) | 15.1 (3.1) | 11 (2) |
| ***Clinical*** | | | | | | |
| Disease duration (y) | 5.4 (5.2) | - | 8 (5.2) | - | 10.4 (4.9) | - |
| MoCA (/30) | 26 (2.4) | 26.8 (2.5) | 25.2 (3.5) | 27.4 (2.2) | 25.1 (5.1) | |
| MCI (Y/N) | 25/52 | - | 16/29 | - | 16/28 | - |
| MMSE (/30) | 28.7 (1.2) | 29.4 (1) | 28.2 (2.3) | 29 (1.5) | 28.3 (1.9) | |
| UPDRS-II | 6.6 (4.7) | - | 10.8 (5.7) | - | 9.6 (6.6) | - |
| UPDRS-III | 15.5 (11) | - | 20.5 (12.1) | - | 19.5 (13.1) | - |
| ***Medication*** | | | | | | |
| LEDD (mg/day) | 676 (466) | - | 707 (445) | - | 633 (386) | - |
| ***Neuropsychiatric tests*** | | | | | | |
| BAI (/63) | 9.7 (8) | 2.9 (4) | 11.5 (7.5) | 2.3 (2.6) | 9.6 (6.9) | 3.3 (4.9) |
| BDI-II (/63) | 7.9 (4.9) | 2.6 (2.4) | 7.8 (4.7) | 1.8 (1.7) | 6.7 (6.1) | 4 (3.6) |
| AES (/63) | 32.9 (8.4) | 24.1 (5.1) | 31.2 (7.1) | 25.1 (5.7) | 31.7 (8.6) | 27.7 (9) |

**Table S2- Demographic, clinical and main neuropsychiatric characteristics of the study cohort longitudinally expressed as: mean (standard deviation).** y: years, M/F: Male/Female, MoCA: Montreal Cognitive Assessment, MCI (Y/N): Mild Cognitive Impairment (yes/no), UPDRS-III: Unified Parkinson's Disease Rating Scale-motor examination, LEDD: Levodopa Equivalent Daily Dose, BAI: Beck Anxiety Inventory score, BDI-II: Beck Depression Inventory, second edition score, AES: Apathy Evaluation Scale.

| | **Baseline** | | **3 years** | | **5 years** | |
|---|---|---|---|---|---|---|
| | **PD (N=68)** | **HC (N=25)** | **PD (N=42)** | **HC (N=17)** | **PD (N=29)** | **HC (N=1)** |
| ***Demographic*** | | | | | | |
| Age (y) | 66.4 (8.3) | 66.6 (4) | 70.5 (7.9) | 68.9 (6.1) | 71 (7) | 69 |
| Sex (M/F) | 46/22 | 15/10 | 28/14 | 8/9 | 17/12 | 1/0 |
| Education (y) | 14.8 (3.1) | 14.2 (2.9) | 14.8 (3.1) | 13.4 (3.2) | 14.1 (3.1) | 13 |
| ***Clinical*** | | | | | | |
| Disease duration (y) | 5.2 (5.2) | - | 7.5 (4.7) | - | 9.4 (3.8) | - |
| MoCA (/30) | 26 (2.4) | 26.6 (2.7) | 25.2 (3.6) | 27.2 (2.3) | 25.7 (3.6) | |
| MCI (Y/N) | 22/46 | - | 15/27 | - | 7/22 | - |
| UPDRS-III | 14.8 (11.2) | - | 20.1 (12) | - | 17.6 (12.8) | - |
| ***Medication*** | | | | | | |
| LEDD (mg/day) | 652 (465) | - | 667 (436) | - | 558 (343) | - |
| ***Neuropsychiatric tests*** | | | | | | |
| BAI (/63) | 9.9 (7.9) | 2.4 (3.2) | 11.5 (7.4) | 2.2 (2.6) | 10.2 (7) | 1 |
| BDI-II (/63) | 7.7 (4.9) | 2.6 (2.5) | 7.8 (4.8) | 1.9 (1.7) | 7.1 (6.6) | 5 |
| AES (/63) | 33(8.6) | 24.1 (5.1) | 31.4 (7.1) | 25.1 (5.7) | 30.9 (7.6) | 37 |

**Table S3- Affiliation of the EEG channels to the four lobes of interest**

| Lobes | EEG channels |
|---|---|
| **Frontal lobe** | E2, E3, E4, E5, E6, E7, E8, E11, E12, E13, E14, E15, E16, E17, E19, E20, E21, E22, E23, E24, E26, E27, E28, E29, E30, E33, E34, E35, E36, E38, E39, E40, E41, E42, E43, E47, E48, E49, E50, E51, E55, E56, E57, E58, E61, E62, E63, E64, E68, E69, E194, E195, E196, E197, E198, E202, E203, E204, E205, E206, E207, E210, E211, E212, E213, E214, E215, E220, E221, E222, E223, E224. |
| **Parietal lobe** | E9, E44, E45, E52, E53, E59, E60, E65, E66, E70, E71, E72, E74, E75, E76, E77, E78, E79, E80, E81, E84, E85, E86, E87, E88, E89, E90, E96, E97, E98, E99, E100, E101, E109, E110, E119, E128, E129, E130, E131, E132, E140, E141, E142, E143, E144, E152, E153, E154, E155, E161, E162, E163, E164, E170, E171, E172, E173, E179, E180, E181, E182, E183, E184, E185, E186, E192, E193. |
| **Temporal lobe** | E67, E73, E82, E83, E91, E92, E93, E94, E95, E102, E103, E104, E105, E111, E112, E177, E178, E188, E189, E190, E191, E199, E200, E201, E208, E209, E216, E217, E218, E219, E225, E227, E228, E229, E231, E232, E233, E235, E236, E237, E239, E240, E242, E243, E245, E246, E247, E249, E250, E251, E253, E254, E255, E256. |
| **Occipital lobe** | E106, E107, E108, E113, E114, E115, E116, E117, E118, E120, E121, E122, E123, E124, E125, E126, E127, E133, E134, E135, E136, E137, E138, E139, E145, E146, E147, E148, E149, E150, E151, E156, E157, E158, E159, E160, E165, E166, E167, E168, E169, E174, E175, E176, E187. |

**Table S4- Affiliation of brain regions to the seven lobes of interest**

| Lobes of Interest | Brain regions |
|---|---|
| **Prefrontal cortex (PFC)** | Middle frontal gyrus (L/R) |
| | Orbito-frontal cortex (L/R) |
| | Superior frontal gyrus (L/R) |
| | Inferior frontal gyrus (L/R) |
| **Motorstrip (Mot)** | Precentral gyrus (L/R) |
| | Paracentral lobule (L/R) |
| | Postcentral gyrus (L/R) |
| **Parietal lobe (Par)** | Inferior parietal lobule (L/R) |
| | Superior parietal lobule (L/R) |
| | Precuneus (L/R) |
| **Temporal network (Tmp)** | Inferior temporal gyrus (L/R) |
| | Middle temporal gyrus (L/R) |
| | Superior temporal gyrus (L/R) |
| | Parahippocampal gyrus (L/R) |
| | Fusiform gyrus (L/R) |
| | Posterior superior temporal sulcus (L/R) |
| **Occipital lobe (Occ)** | Lateral occipital cortex (L/R) |
| | MedioVentral occipital cortex (L/R) |
| **Limbic lobe (Lmb)** | Caudal cingulate gyrus (L/R) |
| | Ventral cingulate gyrus (L/R) |
| | Dorsal cingulate gyrus (L/R) |
| **Insular lobe (Ins)** | Insular gyrus (L/R) |

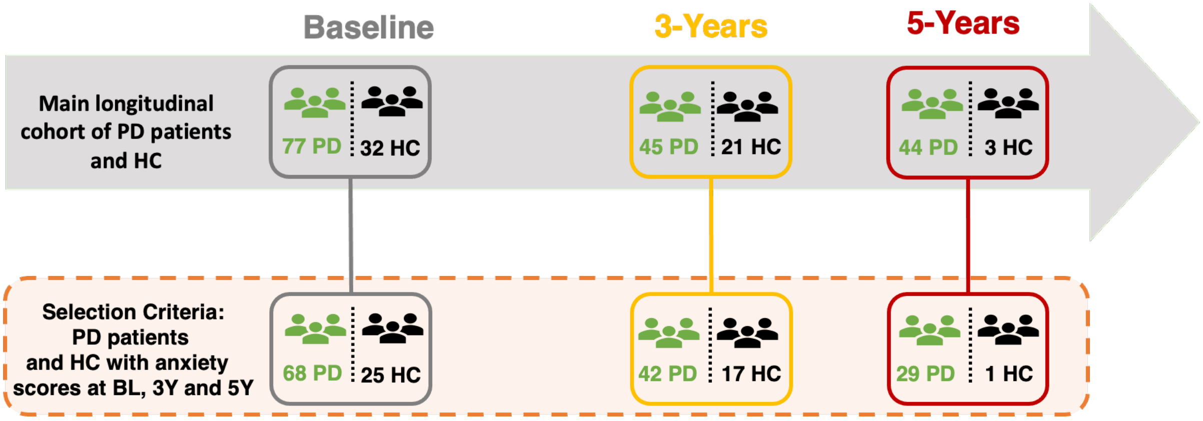


**Figure S1- Flowchart of the main study cohort and the subcohort of participants included in this study.**

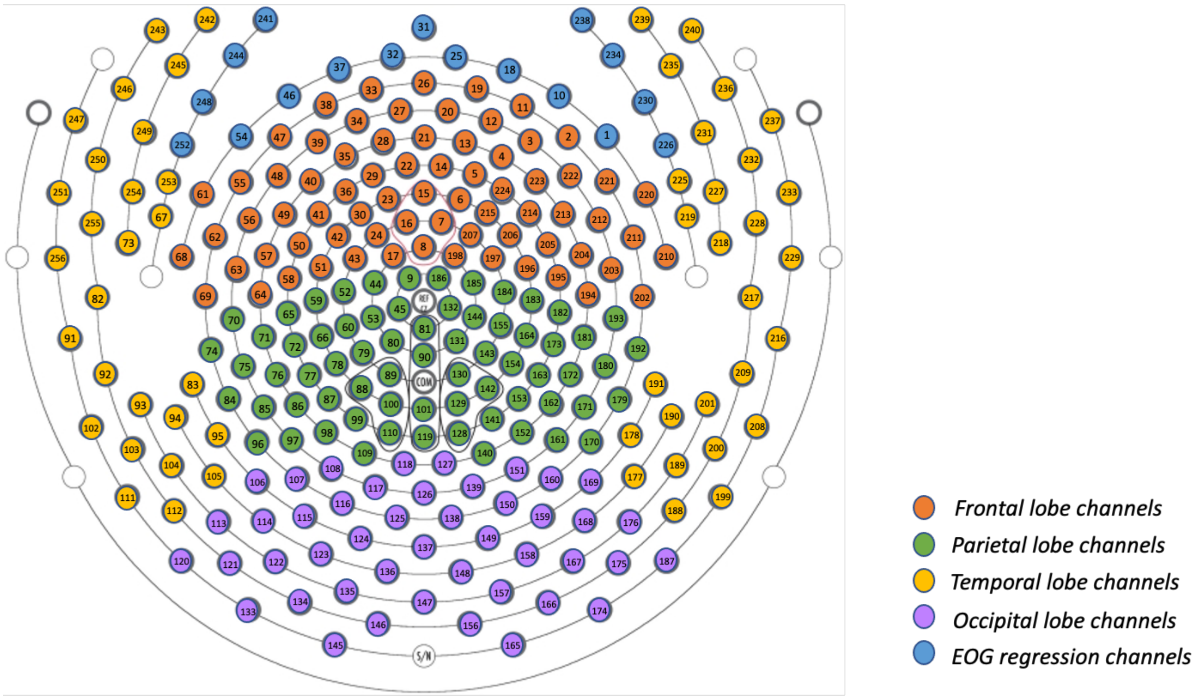


**Figure S2- EGI 256-channels sensor layout and the affiliation of the channels to the four lobes of interest:** Frontal lobe (orange), Parietal lobe (green), temporal lobe (yellow) and occipital lobe (purple). The 17 channels used for EOG regression are marked in blue.
